# A model of inborn metabolism errors associated with adenine amyloid-like fiber formation reduces TDP-43 aggregation and toxicity in yeast

**DOI:** 10.1101/2024.12.03.626668

**Authors:** Sangeun Park, Sei-Kyoung Park, Susan W. Liebman

**Author notes:** Contact: Susan W. Liebman, Department of Pharmacology, University of Nevada, 1664 N Virginia, Mail stop 318, Reno, NV 89557, Lab.

## Abstract

TDP-43 is linked to human diseases such as amyotrophic lateral sclerosis (ALS) and frontotemporal degeneration (FTD). Expression of TDP-43 in yeast is known to be toxic, cause cells to elongate, form liquid-like aggregates, and inhibit autophagy and TOROID formation. Here, we used the *apt1Δ aah1Δ* yeast model of disorders of inborn errors of metabolism, previously shown to lead to intracellular adenine accumulation and adenine amyloid-like fiber formation, to explore interactions with TDP-43. Results show that the double deletion shifts the TDP-43 aggregates from a liquid-like, toward a more amyloid-like, state. At the same time the deletions reduce TDP-43’s effects on toxicity, cell morphology, autophagy, and TOROID formation without affecting the level of TDP-43. This suggests that the liquid-like and not amyloid-like TDP-43 aggregates are responsible for the deleterious effects in yeast. How the *apt1Δ aah1Δ* deletions alter TDP-43 aggregate formation is not clear. Possibly, it results from adenine/TDP-43 fiber interactions as seen for other heterologous fibers. The work offers new insights into the potential interactions between metabolite-based amyloids and pathological protein aggregates, with broad implications for understanding protein misfolding diseases.

## Introduction

Certain proteins and peptides have been shown to be associated with disease when they form amyloid (or amyloid-like) oligomers or fibers. Amyloids are ordered protein aggregates characterized by a cross-β sheet structure that binds to thioflavin T [1]. Such proteins include: Aβ, α-synuclein, TDP-43 (TAR DNA-binding protein 43), FUS (fused in sarcoma), Huntingtin, and p53 which each form aggregates associated, respectively, with Alzheimer’s, Parkinson’s, amyotrophic lateral sclerosis (ALS)/ frontotemporal degeneration (FTD), Huntington’s, and cancer [2–4]. Many of these proteins can also form liquid-like droplets through liquid–liquid phase separation in the cytoplasm. These droplets can be dissolved by hexanediol, exhibit dynamic properties, and can convert into amyloid fibrils [5].

In most cases it is unclear which aggregate form is toxic. Indeed, the very structural nature of TDP-43 aggregates, whether found in patients, or formed in vitro, remains a subject of debate [6]. Some studies suggest that the low-complexity C-terminal domain of TDP-43 forms amyloid in vitro [7], while others report that neither in vitro-formed nor neuron-based TDP-43 aggregates exhibit amyloid properties [6, 8]. Although TDP-43 aggregates are linked to disease, it is unclear whether they contribute to disease pathology or are merely a consequence of it.

Expression of TDP-43 in yeast causes toxicity [9] and a dramatic elongation of cell shape [10]. Mutations in TDP-43 which either increase or decrease its toxicity in yeast, hint that amyloid aggregates may actually have a protective role. Toxicity-enhancing intragenic mutations reduced TDP-43 hydrophobicity, while toxicity-reducing mutations increased hydrophobicity and encouraged larger TDP-43 aggregates in the cytoplasm [11]. However we found that expression of both wild-type and non-toxic TDP-43 protein in yeast formed aggregates that were dissolved by hexanediol and were not stained with thioflavin T [10]. This indicates that TDP-43 forms liquid-like droplets and not amyloid in yeast.

Expression of FUS in yeast is also toxic. But unlike TDP-43, FUS aggregates stain with thioflavin T and do not disappear when cells are treated with hexanediol [10, 12–14].

Extragenic mutations also affect toxicity when TDP-43 is expressed in yeast [9, 15–20]. The mechanism by which TDP-43 causes toxicity remains unknown. However, we found that expression of wild-type TDP-43 in yeast reduces autophagy and TOROID (TORC1 organized in inhibited domain) formation, and extragenic and intragenic mutations which alleviate TDP-43 toxicity, reverse this effect [10, 20]. TOROIDs are large, helical structures of TORC1 that form near the vacuole and are associated with reduced autophagy inhibition.

Yeast prions also form amyloid. [*PSI*^+^], is the prion form of the translational release factor, SUP35. We showed that the de novo appearance of [*PSI*^+^] is dramatically enhanced by the presence of another yeast prion, [*PIN*^+^] (the prion form of RNQ1). Both the [*PSI*^+^] and [*PIN*^+^] prions can form different heritable amyloid shapes with distinct characteristics, called prion strains. Some [*PIN*^+^] strains destabilize [*PSI*^+^] strains and different [*PIN*^+^] strains preferentially promote the appearance of different [*PSI*^+^] strains . Other heterologous amyloid aggregates can also facilitate the de novo appearance of [*PSI*^+^] [21–27].

This type of interaction also occurs in mammalian cells. Aggregates of α-synuclein facilitate fibrillization of tau [28], and Aβ polymerization is promoted by misfolded type-2 diabetes islet amyloid polypeptide [29, 30]. This suggests that cross-seeding is a risk factor for disease.

Recently individual amino acids, and non-proteinaceous metabolites including adenine have also been shown to form amyloid-like fibers that bind ThT and are toxic. Since inborn errors of metabolism disorders lead to accumulation of these elements and cause neurological symptoms a “generic amyloid hypothesis” has been proposed to include protein misfolding diseases as well as inborn errors of metabolism diseases [31–33].

Since these diseases are associated with an increased frequency of neurodegeneration and cancer, it is possible that the metabolite fibers serve as scaffolds upon which pathological protein aggregation can initiate [34].

A model for adenine self-assembly in living yeast was established by deleting two enzymes in the adenine salvage pathway: *APT1* (encoding adenine phosphoribosyltransferase) and *AAH1* (encoding adenosine deaminase). This double deletion causes a dramatic increase in the level of intracellular adenine which increases further when cells are grown in the presence of adenine. Furthermore, the increased intracellular adenine is amyloid-like and associated with growth inhibition [32].

Here, we use this yeast model to show that *apt1Δ aah1Δ* double deletions alter the properties of human TDP-43 expressed in yeast.

## Results

### Examining toxicity of TDP-43 in yeast in the presence of *apt1Δ aah1Δ*

As reported previously when TDP-43-YFP is expressed in yeast on galactose it forms foci, reduces yeast growth rate, and causes an elongation of cell shape [9, 35]. The double *apt1Δ aah1Δ* deletion (ΔΔ) significantly reduces this toxicity (Fig. 1A) and restores cells to their normal shape (Fig. 1B). The ΔΔ mutant also causes a significant reduction in the number of cells with a multiple vs single TDP-43-YFP foci per cell (Fig. 1B).

**Figure 1.**
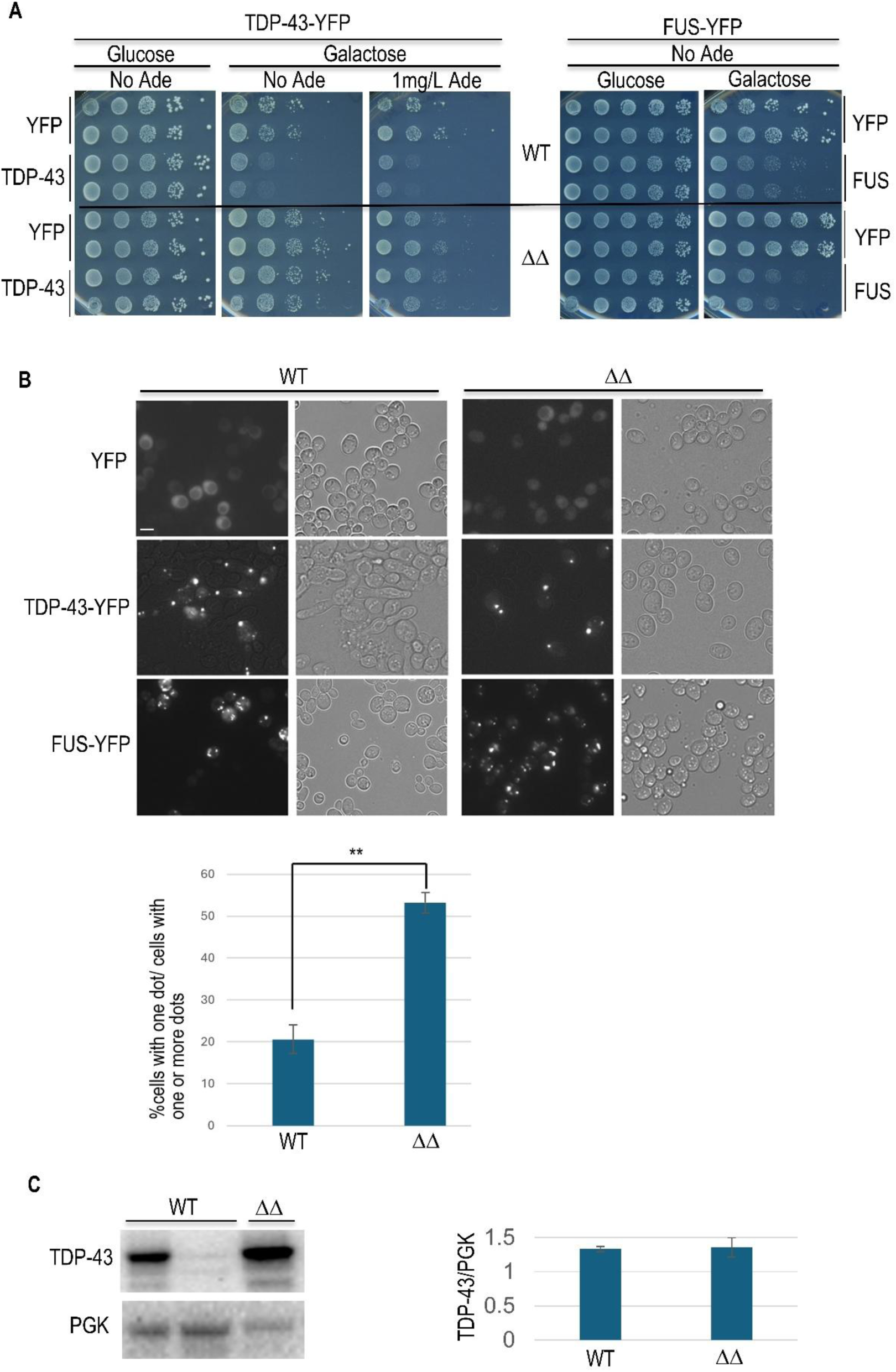
The *aah1Δ apt1Δ* deletions alleviate TDP-43 but not FUS toxicity. (A) The double *aah1*Δ *apt1*Δ deletion alleviated toxicity associated with expression of TDP-43. TDP-43-YFP or FUS-YFP were expressed from the *GAL* promoter on *CEN* plasmids (*YCpGAL-TDP43-YFP*, and *YCpGAL-FUS-YFP*) in wild-type BY4741 [*PIN*^+^] (WT, upper) or an isogenic double *aah1*Δ *apt1*Δ deletion strain (ΔΔ, lower). Transformants were serially diluted and spotted on plasmid selective glucose or galactose plates where the expression of vector-encoded protein was induced. YFP-expressing cells were used as controls (*YCpGAL-EYFP*). No relief of FUS toxicity was observed in the *aah1*Δ *apt1*Δ strain compared to the wild type. (B) The double *aah1*Δ *apt1*Δ deletion changes the appearance of TDP-43 foci and alleviates the elongated cell shape caused by expression of TDP-43. Cells expressing YFP, TDP-43-YFP, or FUS-YFP on selective galactose plates were collected and examined for YFP fluorescence (left) and bright-field microscopy (right). All images were taken at the same magnification. Scale bar, 10 µm. Graph below shows the percentage of cells containing a single TDP-43-YFP aggregates vs. total cells with one or more aggregate. Data are presented as the mean ± standard error of the mean (n=3). More than 500 cells were counted. Statistical significance was determined with a two-tailed t-test (** = p < 0.005). (C) The double deletion doesn’t reduce TDP-43 expression. Left shows a representative Western blot of the WT and double deletion (ΔΔ) strains with galactose induced TDP-43 (*pGAL-TDP43-L*) or an empty vector (*pGAL-L*) with loading control phosphoglycerate kinase (PGK). The lane in the middle was a control without TDP-43. The bar chart includes data from three independently made ΔΔ strains each run on three separate gels. Error bars represent the SEM.

A reduction in TDP-43 toxicity is observed with or without the addition of adenine to the media (Fig. 1A) and this is not caused by a reduction in TDP-43 expression (Fig. 1C). Addition of adenine doesn’t affect the growth rate of wild-type cells but as reported previously [32] it reduces the growth rate of cells bearing the *apt1Δ aah1Δ* double deletion (Fig. 1A). The double deletion was specific for TDP-43, it didn’t alter the toxicity of FUS (Fig. 1A).

### Examining the effect of *apt1Δ aah1Δ* on inhibition of autophagy and TOROID formation caused by TDP-43 expression

The expression of wild-type TDP-43 in yeast reduces autophagy relative to the vector control [20]. As in our previous papers, we measured autophagy by using immunoblotting to examine the breakdown of GFP-ATG8 (Green Fluorescent Protein fused to autophagy-related protein 8) (Fig. 2A) and by examining the cellular location of GFP in cells expressing GFP-ATG8 (Fig. 2B) [10, 20]. We previously found that several modifiers that reduce toxicity of TDP-43 reduce its inhibition of autophagy [10, 20]. Similarly, our current results using both immunoblotting and cellular location measures of autophagy show that the level of autophagy is not altered by *apt1Δ aah1Δ* in the absence of TDP-43. However, both methods show that the presence of *apt1Δ aah1Δ* prevented overexpression of TDP-43 from inhibiting autophagy (Fig. 2AB).

**Figure 2.**
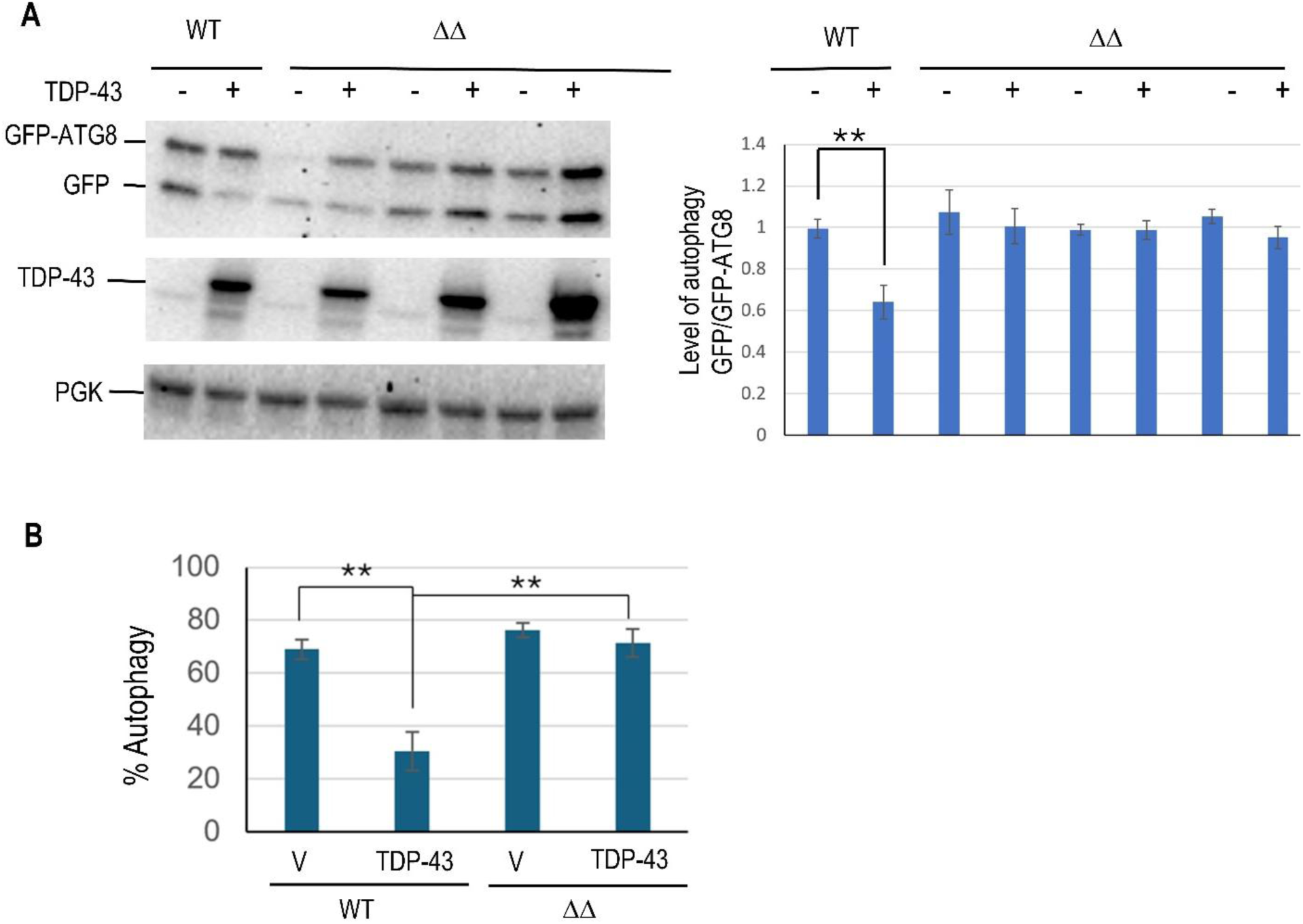
The *aah1Δ apt1Δ* double deletion reverses the inhibition of autophagy caused by overexpression TDP-43. (A) Autophagy measured by Western blots. Autophagy measured by the breakdown of GFP-ATG8 is shown on Western blots. BY4741 [*PIN*^+^] (WT) and three independently made isogenic double deletion strains (ΔΔ) were transformed with *pGAL-TDP43-L* and p*CUP1-GFP-ATG8* and grown on 2% galactose plates with 1% raffinose, the required amino acids methionine and histidine, and 50 µM copper sulfate for 16 hrs at 30°C before being harvested for Western blotting. One representative blot is shown on the left. The bar chart shows data from three independent double deletion strains repeated three time each. (B) Autophagy measured microscopically. Double transformants in BY4741 [*PIN*^+^] (WT) and isogenic double deletion strains shown in (A) were assayed for autophagy by determining the cellular location of GFP-ATG8 microscopically. Autophagy was measured as the fraction of cells with either no GFP-ATG8 fluorescence or with fluorescence only in the vacuole over total live cells. Error bars represent the SEMs calculated from average of three independent transformants by examining 250-550 cells per transformants. ** indicates p <0.005 in a paired two-tailed t-test.

Previously described modifiers that reduce TDP-43’s inhibition of autophagy were also found to lessen TDP-43’s inhibition of TOROID formation, measured by the appearance of TOR1-GFP foci [10]. Likewise we find that *apt1Δ aah1Δ* partially restores TOROID formation in cells expressing TDP-43 (Fig. 3).

**Figure 3.**
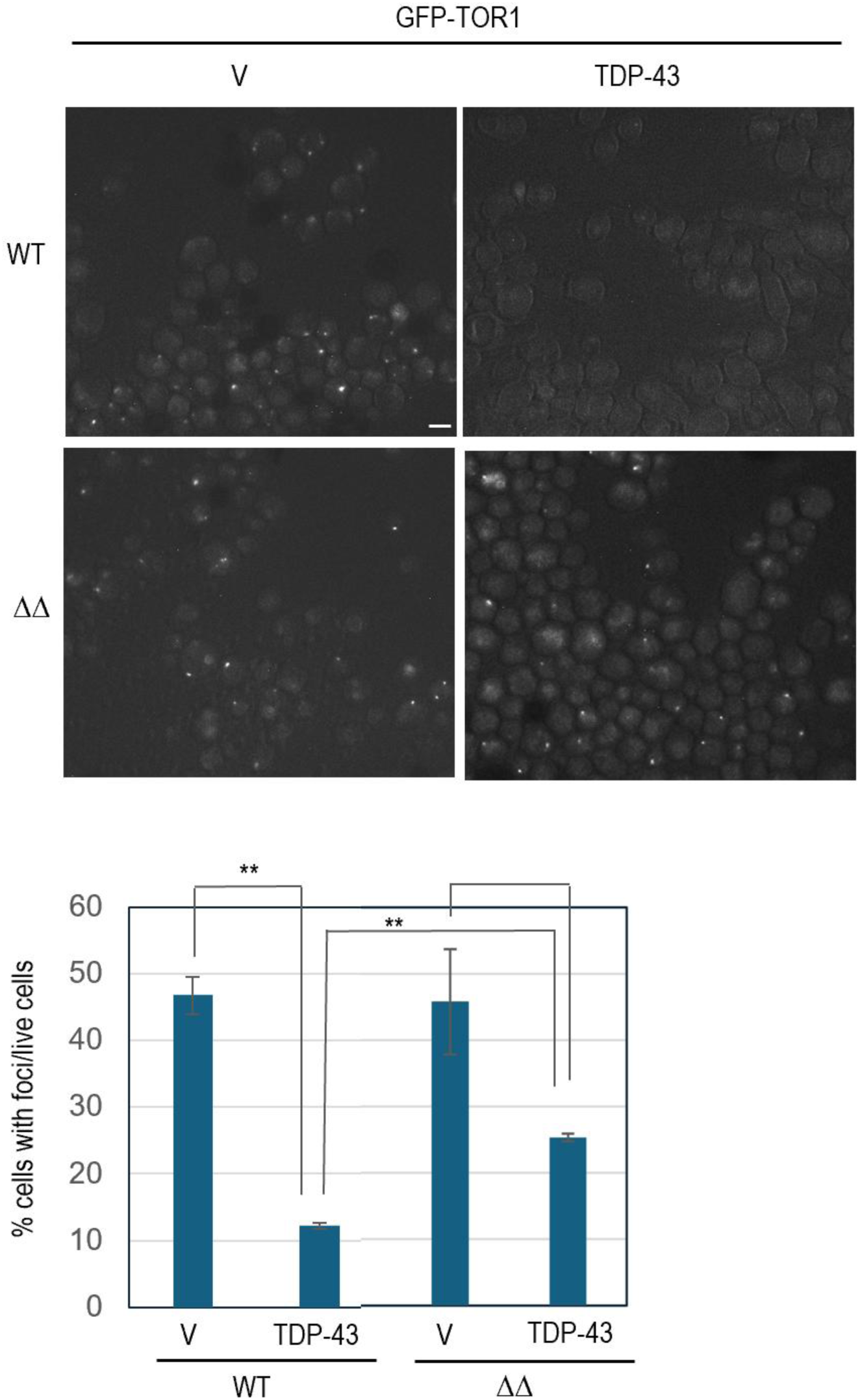
The *aah1Δ apt1Δ* double deletion reduces the inhibition of TOROID formation caused by TDP-43 overexpression. Shown are fluorescent images of endogenously tagged *TOR1* (*GFP-TOR1*) [*PIN*^+^] BY4741 cells with (ΔΔ) or without (WT) the double *aah1Δ apt1 Δ* deletion. Cells were transformed with untagged TDP-43 expressed under the *GAL* promoter in *YCpGAL-TDP43-U* (TDP-43) or empty vector (*YCpGAL-U*, v), grown on selective galactose media for 2 days, examined with a FITC filter and photographed (left). Size bar is 10 µm. The percentage of cells with a GFP-TOR1 dot (TOROID) among live cells expressing TDP-43 or empty vector was counted after 0.5% trypan blue staining to exclude dead cells. Live cells with GFP-TOR1 cytoplasmic foci were averaged from 3 independent transformants by examining about 500 cells per transformant. Error bars are SEMs from three independent transformants. Statistical analysis (paired two-tailed t-test) revealed the following p-values respectively: BY (WT) vector vs. BY (WT) TDP-43 0.000258519, ΔΔ (*aah1Δ apt1Δ*) vector vs. ΔΔ TDP-43 0.062186054, BY (WT) TDP-43 vs. ΔΔ TDP-43 6.41005E-05. Asterisks (**) indicate p < 0.005 in paired two-tailed t-tests. The absence of asterisks denotes no significant difference.

### Examining the effect of *apt1Δ aah1Δ* on the nature of TDP-43 foci

In wild-type strains TDP-43-YPF usually forms one large irregularly shaped focus as well as multiple small foci. In these cells no foci are detected with thioflavin T staining and the TDP-43-YFP foci were all dissolved by hexanediol indicating that they are liquid-like rather than amyloid. In contrast, foci of FUS-YFP stain with thioflavin T and were not dissolved by hexanediol, suggesting their amyloid nature [10].

In an isogenic strain with *apt1Δ aah1Δ* the frequency of multiple small TDP-43-YFP foci is reduced and the larger foci are more punctate than in the wild-type (Fig. 1B). Furthermore, some of these foci stain with thioflavin T (Fig. 4A). While these TDP-43-YFP thioflavin T foci are not as bright as seen for FUS-YFP foci, they are clearly visible. Also, while many of the thioflavin T foci that appear in the cells expressing TDP-43-YFP are also visible in the YFP channel, some are not. Apparently, other proteins, as well as the TDP-43-YFP, have become more amyloid like in the *apt1Δ aah1Δ* strain. Indeed, thioflavin T foci are seen in *apt1Δ aah1Δ* but are never seen in wild-type strains not expressing either TDP-43 or FUS (Fig. 4B).

**Figure 4.**
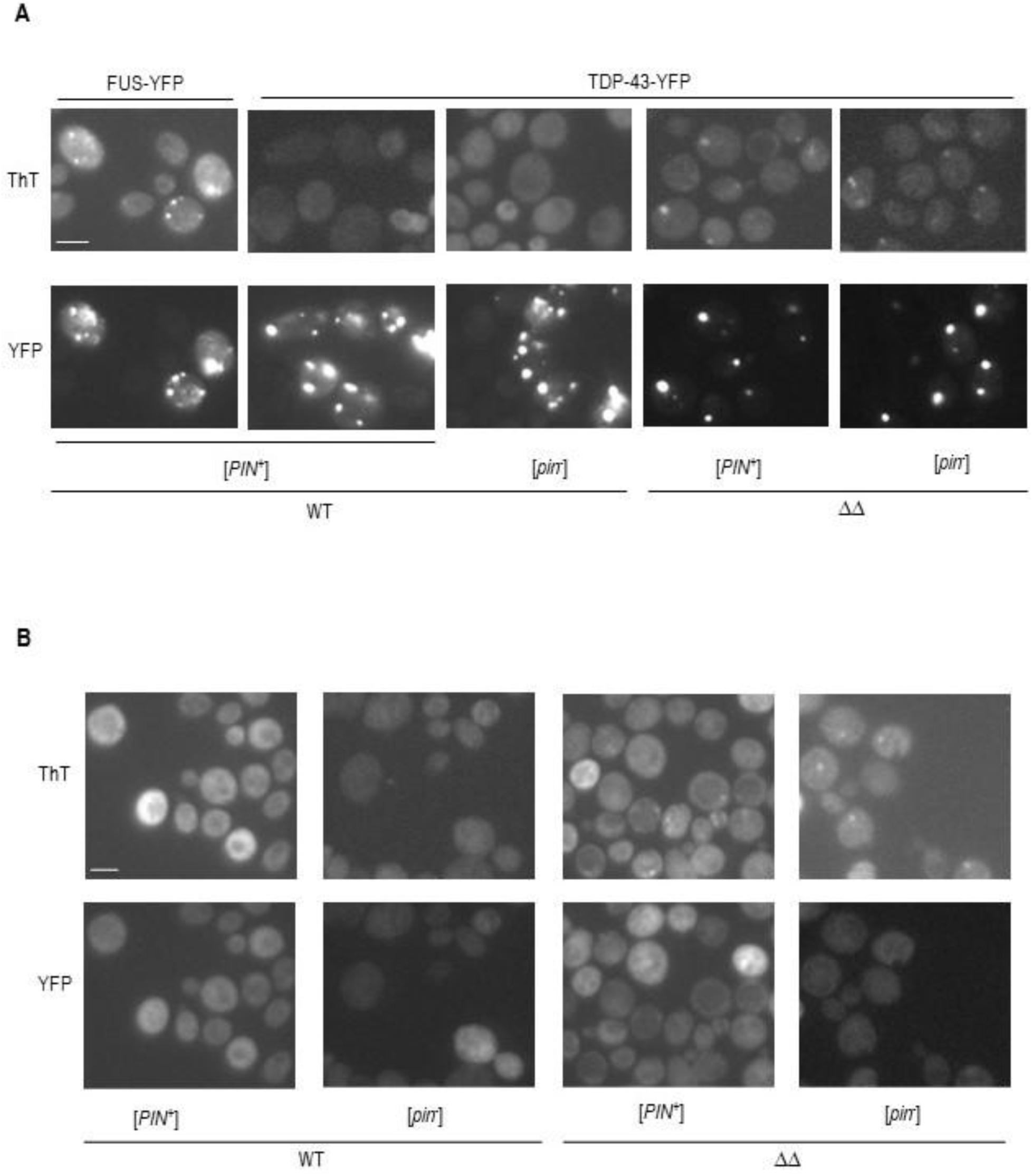
The double *aah1Δ apt1 Δ* deletion makes TDP-43 aggregates more amyloid-like. (A) TDP-43 forms thioflavin-T foci in *aah1Δ apt1Δ* deletion strains. [*PIN*^+^] or [*pin*^-^] BY4741(WT) and isogenic double deletion strains (ΔΔ) were transformed with YC*pGAL-TDP43-YFP-U* or *YCpGAL-FUS-YFP-U*. Cells grown on selective galactose plates lacking adenine and expressing TDP-43-YFP or FUS-YFP were collected, stained with Thioflavin-T (ThT) and examined for CFP fluorescence to see the ThT staining (Top), and YFP fluorescence (Bottom). (B) Thioflavin-T foci formed in *aah1Δ apt1Δ* deletion strains in the absence of TDP-43. [*PIN*^+^] or [*pin*^-^] isogenic wild-type (WT) and *aah1Δ apt1 Δ* deletion strains were transformed with *pYCpGAL-YFP-U*. Cells expressing YFP on selective galactose plates lacking adenine were collected, stained, and examined as in (A). Scale bar is 10 µm.

At the same time as the *apt1Δ aah1Δ* double deletion makes TDP-43-YFP more amyloid-like it reduces TDP-43-YFP’s liquid-like nature. In *apt1Δ aah1Δ,* but not wild-type strains, many TDP-43-YFP foci remain undissolved following hexanediol treatment (Fig. 5).

**Figure 5.**
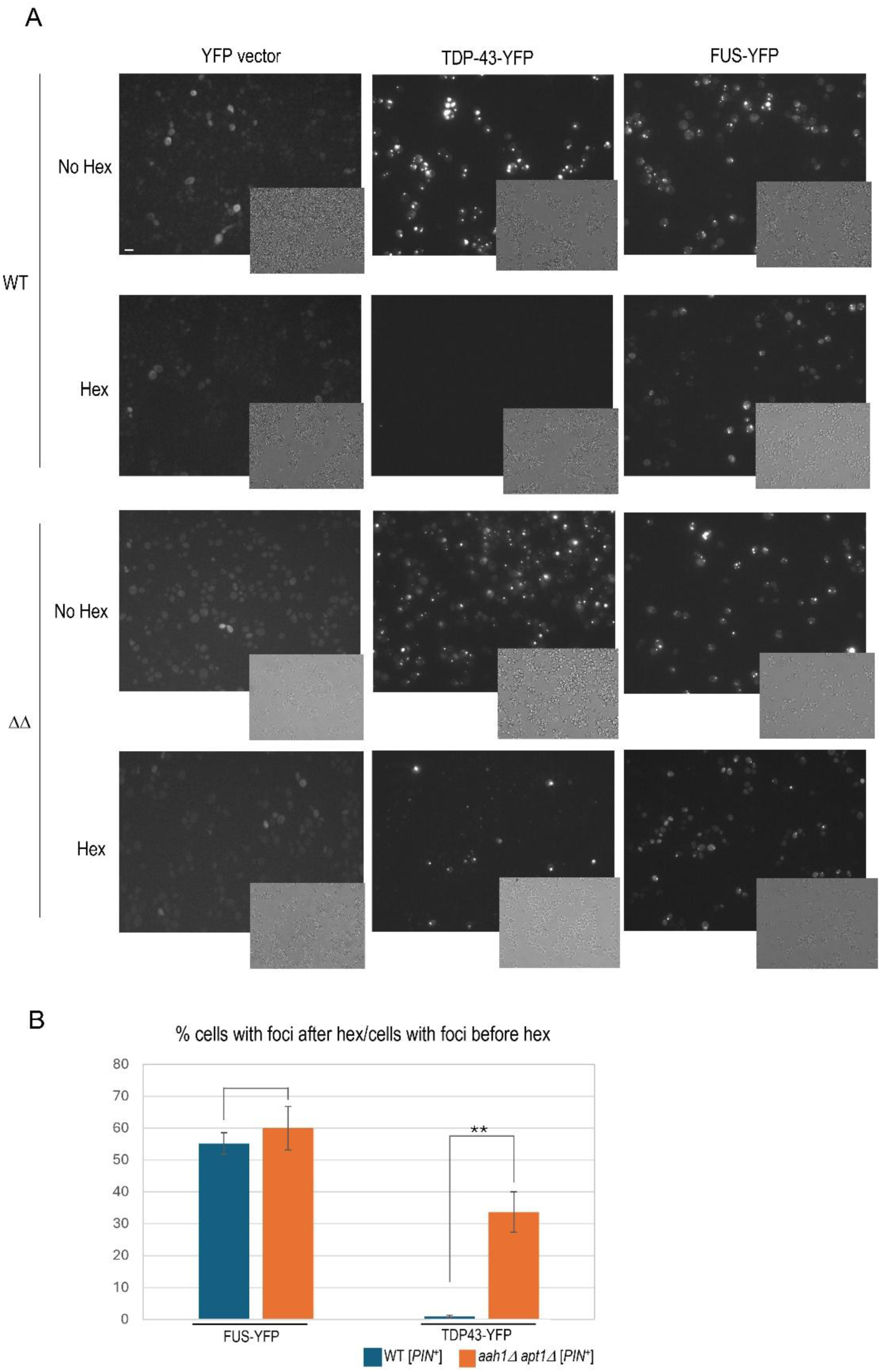
The double *aah1Δ apt1 Δ* deletion strain makes TDP-43 aggregates less liquid-like. Isogenic wild-type (WT) and *aah1Δ apt1 Δ* deletion strains transformed with plasmid expressing YPF (YFP vector), TDP-43-YFP or FUS-YFP were grown overnight in plasmid selective galactose media lacking adenine. Their YFP fluorescence was examined prior to (No Hex) and following treatment with 10% 1,6-hexanadiol for 5 min that dissolves liquid-like aggregates. Three transformants each were tested and gave similar results. (A) Cell under microscope. Fluorescent pictures were taken with 40ms exposure. Bright field images of cells are in inset. Scale bar is 10 µm. (B) Bar Chart. ** indicates p <0.005 in a paired two-tailed t-test. Absence of stars on left indicates no statistical difference.

## Discussion

Previous studies demonstrated the ability of amyloids to influence the aggregation of heterologous proteins, either promoting or inhibiting amyloid formation and by changing the type (strain) of amyloid promoted [21–27]. Here, we sought to determine whether adenine fibers could affect the aggregation and toxicity of TDP-43 in yeast.

The results suggest that the accumulation of adenine, and by extension, the formation of adenine amyloid-like fibers, has a significant impact on the aggregation properties and toxicity of TDP-43 in yeast. The double deletion of *APT1* and *AAH1,* previously shown [32] to cause the accumulation of adenine amyloid-like fibers, reduces the toxicity of TDP-43 and the ability of TDP-43 to elongate cells. This occurs without lowering the levels of TDP-43 in the cell. Notably, these effects are not further enhanced by increasing adenine levels, which is surprising given that elevated adenine increases amyloid-like adenine fiber formation [32]. This lack of additional effect may be due to adenine itself being toxic at higher concentrations in the double deletion strain, thus masking any beneficial impact it might have on TDP-43 toxicity. As seen for other modifiers of TDP-43 toxicity [10, 20], the double deletion reduces the inhibition of autophagy and TOROID formation caused by expression of TDP-43. This occurs even though the double deletion itself has no effect on autophagy or TOROID formation in the absence of TDP-43. Strikingly, the double deletion causes TDP-43 aggregates to become less-liquid like and more amyloid like, suggesting that amyloid-like TDP-43 foci are less toxic than liquid-like TDP-43 foci and less able to disrupt autophagy and TOROID formation.

One possible explanation for these effects is cross-seeding between adenine fibers and TDP-43. Cross-seeding is a well-established phenomenon where aggregates of one amyloid protein can promote or inhibit the aggregation of another. In vitro studies have shown that hydrogels of hnRNPA1 can directly cross-seed the aggregation of FUS, another amyloid-forming protein [36]. It is plausible that a similar mechanism is at play here, where adenine fibers interact with TDP-43 and alter its aggregation state, thereby reducing its toxicity.

However, this hypothesis raises an interesting question: if adenine fibers are indeed interacting with TDP-43, why does the double deletion reduce TDP-43 toxicity even without added adenine? This could be explained by the fact that the double deletion leads to a baseline induction of intracellular adenine amyloid-like fiber even in the absence of exogenous adenine [32]. Possibly this low level of fibers is sufficient to saturate the effect on TDP-43 aggregation, such that any additional adenine beyond this threshold level does not further influence the properties of TDP-43 aggregates.

The ability of excess cellular adenine to influence TDP-43 aggregation properties is consistent with the idea that amyloid-like adenine fibers can affect other cellular amyloids or aggregates. This suggests a broad role for metabolite-based amyloid structures in modulating the behavior of pathological protein aggregates in cells.

An alternative explanation for the observed effects could be indirect. Rather than direct cross-seeding between adenine fibers and TDP-43, the double deletion may be affecting TDP-43 toxicity through changes in other cellular pathways. For example, the accumulation of adenine could be altering the levels of molecular chaperones or other components of the protein quality control machinery, which in turn could modulate the aggregation and toxicity of TDP-43.

## Conclusion

In summary, our findings suggest that the double deletion of *APT1* and *AAH1* reduces TDP-43 effects on toxicity, autophagy, and TOROID formation by altering the properties of its aggregates, potentially through cross-seeding with adenine fibers. The fact that this effect occurs even in the absence of added adenine points to a threshold level of adenine amyloid-like fiber formation that is sufficient to impact TDP-43. These results open up new avenues for exploring the role of metabolite-based amyloids in modulating protein aggregation and toxicity, with potential implications for understanding the broader landscape of protein misfolding diseases. Further studies will be required to determine whether these effects are mediated through direct interactions between adenine fibers and TDP-43 or through indirect mechanisms involving changes in cellular protein homeostasis.

## Materials and Methods

### Strains and plasmids

Strains used are shown in Table 1. To construct the *aah1Δapt1Δ* strain, we began with BY4741[37] containing *apt1:: hphMX6* obtained from Matin Kupiac’s collection via Dana Laor [32] and BY4741 *aah1Δ* from the deletion library [38]. We transformed the BY4741 *apt1:: hphMX6* strain with a PCR-amplified *aah1::KanMX* fragment from the other strain using upstream and downstream gene-specific 45mer primers. Cells were first grown on a YPD plate before being plated on YPD containing 200 μg/ml G418 to select for double deletion strains. These strains were further validated by growth on YPD containing 200 μg/ml hygromycin. The gene disruptions were confirmed with PCR.

**Table 1.**
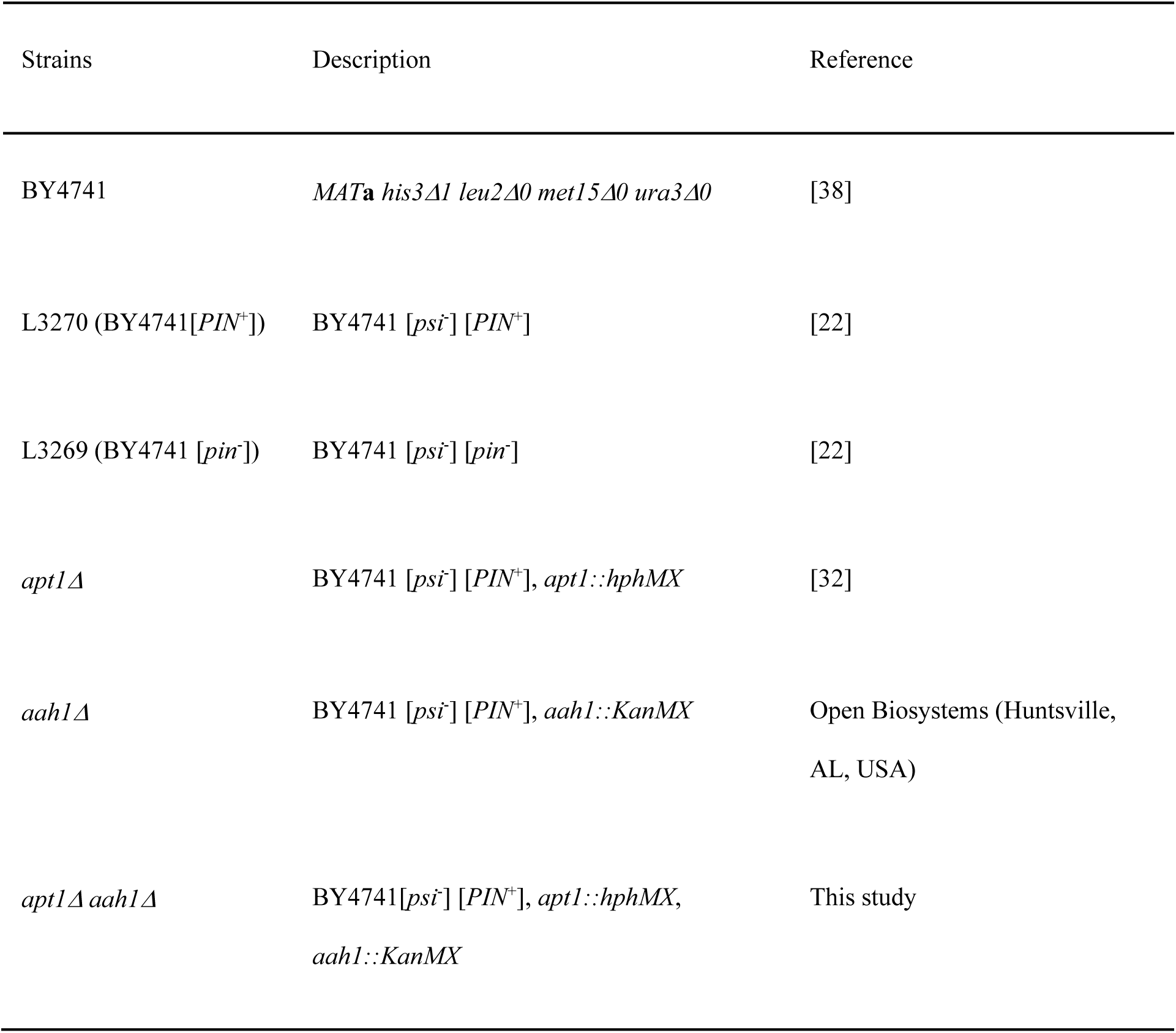
Strains.

To measure TOROID formation, we disrupted *TOR1* while integrating *GFP-TOR1* controlled by its native promoter into BY4741. Wild-type BY4741 or *aah1Δapt1Δ* strains were transformed with SpeI-digested integrating pSK108 plasmid containing the TOR1 promoter and GFP tagged N-terminal region (kindly sent by Dr. Noda)[39] and selecting for transformants on leucineless media. Spel makes a unique cut in the N-terminal *TOR1* region on the plasmid. The correct insertions were confirmed with PCR. Plasmid pSK108 is shown in Table 2.

**Table 2.** Plasmids.

| Short name, SWL laboratory plasmid # | Description | Reference |
| --- | --- | --- |
| pSK108 | pRS305--PTOR1 (1000 bases)-GFP-TOR1 (426 N-terminal bases), LEU2 | [39] |
| YCpGAL-EYFP, p2260 | pAG416 Gal-ccdB-EYFP (*URA3*, *CEN*) | Addgene plasmid #14219, made by Aaron Gitler |
| YCpGAL-TDP43YFP, p2042 | pRS416 Gal-TDP YCpGAL-TDP-43-YFP, (*URA3*, *CEN*) | Addgene plasmid #27447, deposited by Aaron Gitler |
| YCpGAL-FUS-YFP, p2043 | pRS416 Gal-FUS-EYFP (*URA3*, *CEN*) | [14] |
| YCpGAL-U, p2129 | pAG416 Gal-ccdB (*URA3*, *CEN*) | Addgene plasmid #14147, made by Aaron Gitler |
| YCpGAL-TDP43-U, p2665 | pAG416 Gal-TDP-43 (*URA3*, *CEN*) | [10] |
| YCpGAL-L, p2245 | pAG415 Gal-ccdB (*LEU2*, *CEN*) | Addgene plasmid #14145, made by Aaron Gitler |
| YCpGAL-TDP43-L, p2368 | pAG415 Gal-TDP-43 (*LEU2*, *CEN*) | [10] |
| YCpCUP1-GFP-ATG8, p2571 | pCUP1-GFP-ATG8 (*URA3*, *CEN*) | Addgene #49423, deposited by Daniel Klionsky |

### Scoring for Growth

Standard yeast medium was used throughout [40]. To assess growth of cells expressing plasmids, transformants were grown overnight on plasmid selective synthetic glucose medium lacking adenine. They were then normalized to OD_600_ = 4, serially diluted in water 10X, and spotted on synthetic medium with 2% glucose or 2% galactose supplemented with required amino acids, nucleobases, and 0 or 1mg/L adenine. Plates were incubated at 30°C for four days.

### Microscopy for YFP tagged proteins before and after 1,6 Hexanediol treatment, and for Thioflavin T-stained protein

A Nikon Eclipse E600 fluorescent microscope (Nikon, Tokyo, Japan) with 100×/1.23 NA or 60×/1.4 NA oil immersion objectives was used to observe YFP fluorescence, with or without prior 1,6 Hexanediol treatment. Cells were also observed in bright field. Thioflavin T-stained fluorescent foci were observed using a CFP filter. Post-acquisition image processing (brightness and contrast) was performed using Adobe Photoshop 2024 (San Jose, California), and all micrographs display a scale bar. Detailed methods have been described previously [10].

### Scoring for autophagy by determination of GFP-ATG8 cleavage with western blots and microscopic fluorescence counting

BY4741 WT and its isogenic double *aah1*Δ *apt1*Δ deletion strains were doubly transformed with YC*pGAL-TDP-43-L* and YC*pCUP1-GFP-ATG8*. The transformants grown on plasmid selective glucose plates lacking adenine were normalized to 0.2 OD_600_ and spread on TDP-43 and GFP-ATG8 overexpressing plasmid selective galactose plates with 1% raffinose and 50 µM CuSO4. Preparation of lysates and immunoblotting were as described previously [20]. α-GFP (1:5000, Roche); α-TDP-43 (1:3000; Proteintech Group); and α-PGK (yeast 3-phosphoglycerate kinase, 1:10,000, Novex) were respectively used to compare the levels of uncleaved GFP-ATG8 and cleaved GFP, TDP-43, and PGK (yeast 3-phosphoglycerate kinase). The ratio of cleaved GFP to uncleaved GFP-ATG8 measures autophagy. The internal PGK control was used to compare the level of TDP-43 present in wild-type (WT) vs. double deletion strains (ΔΔ).

Microscopic determination of the cellular location of GFP-ATG8 was also used to measure autophagy. Autophagy was calculated as the fraction of cells with either no fluorescence or with fluorescence only in the vacuole over total live cells. Dead cells were detected with trypan blue staining seen in bright field. Microscopic determination of the location of the fluorescence GFP-ATG8 was as previously reported [20].

### Measuring TOROID formation

TOROID formation was measured as described previously [20].

## Acknowledgments

We thank Robbie Joséph Loewith, Daniel J. Klionsky, Aaron Gitller, Susan Lindquist, Martin Kupiac, Dana Laor, and Ehud Gazit for the gifts of plasmids and strains, and Irina Derkatch for the helpful suggestions. This work was funded by a National Institutes of Health MIRA grant (1R35GM136229-04) awarded to S.W.L.

